# Loss of ATP-sensitive channel expression and function decreases opioid sensitivity in a mouse model of type 2 diabetes

**DOI:** 10.1101/2023.09.06.556526

**Authors:** Cole Fisher, Kayla Johnson, Madelyn Moore, Amir Sadrati, Jody L. Janecek, Melanie L. Graham, Amanda H. Klein

## Abstract

During diabetes, β-cell dysfunction due to loss of potassium channels sensitive to ATP, known as K_ATP_ channels occurs progressively over time contributing to hyperglycemia. K_ATP_ channels are additionally present in the central and peripheral nervous systems and are downstream targets of opioid receptor signaling. The aim of this study is to investigate if K_ATP_ channel expression or activity in the nervous system changes in diabetic mice and if morphine antinociception changes in mice fed a high fat diet (HFD) for 16 weeks compared to controls. Mechanical thresholds were also monitored before and after administration of glyburide or nateglinide, K_ATP_ channel antagonists, for four weeks. HFD mice have decreased antinociception to systemic morphine, which is exacerbated after systemic treatment with glyburide or nateglinide. HFD mice also have lower rotarod scores, decreased mobility in an open field test, and lower burrowing behavior compared to their control diet counterparts, which is unaffected by K_ATP_ channel antagonist delivery. Expression of K_ATP_ channel subunits, Kcnj11 (Kir6.2) and Abcc8 (SUR1), were decreased in the peripheral and central nervous system in HFD mice, which is significantly correlated with baseline paw withdrawal thresholds. Upregulation of SUR1 through an adenovirus delivered intrathecally increased morphine antinociception in HFD mice, whereas Kir6.2 upregulation improved morphine antinociception only marginally. Perspective: This article presents the potential link between K_ATP_ channel function and neuropathy during diabetes. There is a need for increased knowledge in how diabetes affects structural and molecular changes in the nervous system to lead to the progression of chronic pain and sensory issues.

## Introduction

ATP-sensitive potassium channels (K_ATP_ channels) are a family of weak inward rectifying potassium channels expressed in diverse cell types including the pancreatic β-cells, heart, brain, skeletal and smooth muscle cells. K_ATP_ channels are most famously involved in glucose stimulated insulin in the pancreas. Glucose metabolism drives an initial rise in ratio of ATP to ADP via oxidative phosphorylation or plasma membrane-associated pyruvate kinase that leads to K_ATP_ channel closure and cell depolarization(1). Coupling cellular energy metabolism to membrane electrical activity is thought to be an important function of these channels across all tissues. K_ATP_ channels are octamers, with four potassium channel subunits, Kir6.1 or Kir6.2, and four sulfonylurea regulatory subunits, either SUR1 or SUR2. Kir6.2-SUR1 is a major subunit combination of K_ATP_ channels not only in the pancreas, but also in vagal, hippocampal, myenteric, spinal cord, and dorsal root ganglia (DRG) neurons.

For several decades, multiple oral medications have been available that promote release of insulin, namely the sulfonylureas and the meglitinides These drug classes, sometimes known as “insulin secretagogues”, block the K_ATP_ channel to depolarize beta cells when glucose and ATP levels are high within cells. Sulfonylurea drug prescribing (e.g. glyburide, glimepiride, gliclazide) has decreased over the years, but are still used in up to 20% of patients in the United States(2). Second generation sulfonylureas offer good glycemic control and are also available at an affordable cost, offering an alternative to newer antidiabetic drugs that is effective in many patients. The mechanism of action of the meglitinides closely resembles that of the sulfonylureas, with two exceptions. The binding site on the sulfonylurea subunit of the K_ATP_ channel is different, and meglitinides have a very short onset of action and a short half-life compared to sulfonylureas, and are best delivered immediately preceding or during a meal to control blood glucose.

Achieving normoglycemia is an important treatment process for patients with T2DM, as hyperglycemia can affect several organ systems, including the nervous system. Diabetic neuropathy is a common nerve dysfunction in people with diabetes, and diagnosis typically involves careful clinical analysis and sensory/motor testing(3). Typically patients with a history of long-standing hyperglycemia and hyperlipidemia and elevated HbA1c are the most at risk for development of neuropathy(4), which affects up to 50% of diabetes patients in their lifetime(5). Treatment options for diabetic neuropathy are incredibly limited, but include tricyclic antidepressants, anticonvulsants (e.g. gabapentin and pregabalin,) and the serotonin and norepinephrine reuptake inhibitors(3). Although opioid drugs are not first-or second-line agents recommended in diabetic neuropathy, they are still common among the treatment options used for this condition(6).

The analgesic effect of opioids, including morphine and other synthetic and semi-synthetic agents, are mediated by μ-, δ-, and κ-opioid receptors expressed in the peripheral and central nervous systems(7). Upon activation by an agonist, opioid receptors couple to G-proteins which dissociate and interact with various intracellular effector systems, including K_ATP_ channels(8; 9). It is likely that multiple signaling pathways targeting ion channels may work to promote hyperexcitability of sensory neurons during neuropathy(10), and medications blocking K_ATP_ channels would be predicted to potentially promote hyperalgesia and/or decrease antinociception of analgesics, particularly morphine. Since several oral antidiabetic drugs target K_ATP_ channels, and opioid medications are still used to treat diabetic neuropathy in the clinic, it would seem that an important drug-drug interaction could be occurring for thousands of patients. We tested the hypothesis that K_ATP_ channel expression would be affected in the spinal cord and dorsal root ganglia (DRG) of diabetic mice and that systemic administration of glyburide and nateglinide would inhibit morphine-induced antinociception in these animals compared to age and sex matched controls.

## Materials and Methods

### Animals

The University of Minnesota Institutional Animal Care and Use Committee (IACUC) approved all animal use procedures discussed herein. Male and female C57BL6 mice (approximately four weeks of age, 13-25 g; Charles River, USA), housed in a standard 12 hr day/night environment, were placed in randomly assigned groups. Beginning at five weeks of age, groups received either a control diet (CON) or a high fat, high carbohydrate diet (HFD, Open Source Diets #12328 and 12330, Research Diets, New Brunswick, NJ, respectively) for 17 weeks. Food and water were available *ad libitum* and staff monitored food and water intake and animal health weekly. The drug and viral treatments were blinded to the experimenter during all behavioral testing.

### Intraperitoneal Glucose Tolerance testing

Animals were fasted for six to eight hours prior to lateral tail vein blood collection. Following completion of a baseline measurement, dextrose was administered (100 µL, 0.5 g/mL dextrose in sterile saline, IP) and blood glucose readings were obtained at 10, 30, and 60 minutes post-dextrose using a glucometer (Alpha Trak2 glucose meter, Zoetis Inc., Kalamazoo, MI).

### Blood assays

Metabolic assays using EMD Millipore’s MILLIPLEX® MAP Mouse Metabolic Hormone Bead Panel (MMHMAG-44k-07, MilliporeSigma, USA) were for: glucose-dependent insulinotropic polypeptide (GIP), C-Peptide 2, and Insulin. Blood was taken using a superficial temporal vein needle prick and was collected in a BD Microtainer® SST™ collection tube (#420075, McKesson Medical-Surgical Inc., USA) prepped with 1-2 µL aprotinin. Animals were given saline (1mL, sc) and ketoprofen (5mg/kg in saline, SC) after blood collection. Blood samples were centrifuged at 3500 rpm for 10 min at 4° C (Eppendorf Centrifuge 5415R). Serum was stored at −80 ° C until processing per manufacturer’s guidelines.

### Mechanical Paw Withdrawal Thresholds

Mice were placed into acrylic animal enclosure cubicles on top of a mesh floor and acclimated for 30-60 minutes a day for one week prior to official testing. Mechanical paw withdrawal thresholds were measured using electronic von Frey testing equipment (Electric von Frey Anesthesiometer, series 2390, Almemo® 2450 Ahlborn, IITC Life Science, Woodland Hills, CA). A probe was pressed into the plantar surface of the hind paw, and the force required to evoke a nocifensive response (e.g. paw lifting, tapping, shaking, licking, jumping or biting at the probe). Five measurements from both the right and left hind paw were obtained for baseline measurements with at least 30 seconds interstimulus period between each measurement.

### Thermal Paw Withdrawal Latency

Mice were placed into clear acrylic cubicles on top of a glass platform heated to 30° C. The Hargreaves method was used to determine thermal withdrawal latency by recording the amount of time a heat radiant beam of light, focused on the plantar surface of the hind paws, was required to cause a nocifensive response (paw lifting, tapping, shaking, jumping or licking the paw; Plantar Test Analgesia Meter, Model 400, IITC, Woodland Hills, CA). The maximum time of exposure was 20 seconds. Five measurements from both the right and left hind paws were obtained with at least 2 minutes between successive tests.

### Rotarod testing

Motor coordination was assessed with the Rotamex-5 automated rotarod system (Model 0254-2002L Columbus Instruments, Columbus, OH) using a 3.0 cm rod assembly. A 4 rpm starting speed was increased in 1 rpm increments at 30-second intervals until animals fell off the rod or reached a top speed of 14 rpm (300 s). Two tests per animal were conducted and averaged.

### Open-Field testing

Animals were placed in an open field arena (40 cm x 40 cm) and activity was recorded with a digital camcorder for offline analysis (Sony Handycam, HDR-CX405; Sony Corp., Tokyo, Japan). Animals were recorded for 15 minutes of baseline activity, followed by injection of morphine (15 mg/kg, sc) and recorded for another 30 minutes. The distance traveled, average velocity, change in angular orientation, and time spent immobile were calculated using Ethowatcher computational software for data analysis (https://ethowatcher.paginas.ufsc.br/).

### Nesting behavior

Animals were placed in individual cages overnight (∼14 hrs) with a ∼5 cm cotton Nestlet^TM^ square (Ancare, Bellmore, NY). In the morning, nest building was assessed by applying the nest complexity scoring matrix(11). Nesting scores were based on a 0-5 scale with 0.5 increments.

### Burrowing

Mice were placed into individual empty cages containing a burrowing tube filled with 500 g of pea gravel for two hours. Tubes were constructed from PVC conduit tubing sections with a 20 cm x 6 cm interior. The open ends of tubes were raised approximately 4 cm with bolts. Any gravel remaining in the tube was weighed to assess gravel displacement.

### Drug Administration

Six groups of animals received daily doses in the morning of either glyburide (50 mg/kg), nateglinide (50 mg/kg), or vehicle (20% DMSO, 0.5% Tween in sterile saline, 100uL) during the last five weeks on either diet (weeks 12-16). Sixteen weeks after starting either a CON or HFD diet, cumulative antinociception dose response curves were obtained for morphine (2.5, 5, and 10 mg/kg, SC in saline). In four additional groups of mice pregabalin (12.5, 25, 50 mg/kg, SC in saline) or gabapentin (25, 50,100 mg/kg, SC in saline) was given 30 minutes after baseline mechanical thresholds were obtained.

### qPCR

RNA was extracted from harvested tissues (spinal cord and dorsal root ganglia) using RNeasy Mini Kit (#74104 QIAGEN, Inc.; Germantown, MD) with DNase I digestion. Standard manufacturers’ guidelines were followed and are described briefly here. A maximum of 30 mg of tissue was macerated in 1 mL TriReagent using a glass tissue grinder before titration with a 22 ga blunt needle. Homogenate was incubated at room temperature for 5 min with 0.1 mL bromo 3-chloropropane before centrifugation. Spinal cord RNA was eluted over two washes of 35 µL RNase free water, whereas dorsal root ganglia RNA was eluted with a single 35 µL wash. RNA was quantified using either Nanodrop (ND-1000, Thermo Fisher Scientific, Wilmington, DE) and/or a Qubit 2.0 Fluorometer (Thermo Fisher Scientific). Reverse transcription with 50 ng of RNA into a cDNA library was performed using the Omniscript RT kit (#205111, QIAGEN, Inc.; Germantown, MD) and random nonamers (Integrated DNA Technologies, Coralville, IA). Samples were amplified in block cycler at 37° C for 60 min, 93° C for 3 min, and held at 4° C.

qPCR was performed using a Roche LightCycler^®^ 480 instrument (Roche Diagnostics, Mannheim, Germany) using LightCycler^®^ 480 SYBR Green I Master mix (Roche Diagnostics, Mannheim, Germany). Temperature-dependent dissociation of the DNA product was examined using a melting curve at the end of each reaction. Data were normalized against an 18s housekeeping gene as used previously (12).

### C-Fiber Compound Action Potentials

Compound action potentials (CAPs) were measured from the sciatic nerves of mice fed either a CON or HFD for 16 weeks. Sciatic nerves were dissected from the hind limbs of mice and recordings were performed the day of harvesting. One nerve from each mouse was mounted in a chamber filled with superficial interstitial fluid. Electrical stimulation was performed at a frequency of 0.3 Hz with electric pulses of 100-μs duration at 100–10,000 μA delivered by a pulse stimulator (2100, AM Systems, Carlsborg, WA, United States). Evoked CAPs were recorded with electrodes placed 10 mm from the stimulating electrodes. Dapsys software was used for data capture and analysis (Brian Turnquist, Bethel University, St. Paul, MN, United States). The stimulus with the lowest voltage producing a detectable response in the nerve was determined as the threshold stimulus. The stimulus voltage where the amplitude of the response no longer increased was determined to be the peak amplitude. The conduction velocity was calculated by dividing the latency period, the time from stimulus application to neuronal initial response, by the stimulus-to-recording electrode distance.

### Adenoviral Upregulation of SUR1 and Kir6.2 subunits

After being fed either a CON or HFD for 15 weeks, six additional groups of mice were inoculated intrathecally with adenovirus (Ad) carrying constructs for either SUR1 (Ad-m-ABCC8, ADV-251792, 3-5e10 PFU, Vector Biolabs, Malvern, PA), Kir6.2 (Ad-m-KCNJ11, ADV-262651, 3-5e10 PFU, Vector Biolabs, Malvern, PA) or a control vector (Ad-CMV-Null, ∼2e10 PFU). For intrathecal injections, a 10 μL inoculum was administered to conscious mice as described previously(13; 14).

### Immunohistochemistry and Microscopy

Seventeen weeks after intrathecal inoculation, posterior hind paw skin was isolated and submerged in Zamboni Fixative (NC9335034, Newcomer Supply, Middleton, Wisconsin) overnight and moved to 1x PBS and placed in 4°C for long term storage. Posterior hind paw skins were embedded in O.C.T., sectioned (10 μM, Leica CM3050) and mounted on electrostatically charged slides. For immunohistochemistry, 1-2 slides from 3 mice per treatment group were incubated at room temperature (RT) in 50 mM glycine for 45 minutes and then washed twice, 5 minutes each, in 1x PBST (0.2% triton x-100). Tissues were then blocked for 1 hr at RT in PBST + 10% horse serum (26050-088; Gibco, Thermo Fisher Scientific, Waltham, MA) + 1% bovine serum albumin (BP1600-100, Thermo Fisher Scientific) and then incubated with the primary antibody (ab108986, Abcam, Cambridge, United Kingdom) at 1:500 dilution in blocking solution overnight at 4°C. Tissue was then washed three times, 5 minutes each in PBST and incubated with the secondary antibody (ab150075, Abcam) at a 1:1000 dilution in blocking solution for 4 hr at room temperature. Tissues were washed three more times for 5 minutes each in PBST and then mounted using DAPI Fluoromount-G (0100-20, Southern Biotech, Birmingham, AL). Slides were imaged using a 20x objective on a Zeiss laser scanning confocal microscope (LSM710, Carl Zeiss, Oberkochen, Germany).

### Fluorescence Quantification

Confocal microscopy files were analyzed using ImageJ/Fiji (https://imagej.net/Fiji). A region of interest (ROI) was drawn on the image to include the epidermis, dermis, and subcutaneous skin layers along with 3 smaller background regions per image. The same ROI and 3 background regions were used for both fluorophore channels. Initial measurements were taken at this time including area, min & max grey value, integrated density, and mean grey value. Rolling ball radius was used for background subtraction. The corrected total fluorescence was calculated using the equation (integrated density-[area of ROI – mean fluorescence of 3 background samples]). A total of 10 images were included for analysis per animal.

### Data Analysis

Data are presented as mean ± SE or as median ± 95% CI. For normally distributed data, statistical comparisons were made by unpaired t-tests, linear regression, one-way ANOVA or two-way ANOVA with between-group differences identified using Tukey or Dunnett post hoc tests for at least three groups’ comparisons. Non-normal data were analyzed using non-parametric statistics. Statistical analyses were performed using either GraphPad 9 or SPSS (IBM, Version 27). Significance was defined as *P* < 0.05.

## Results

### Development of type 2 diabetic phenotype in mice

Male and female mice were placed on either a control (CON) or high fat diet (HFD) for sixteen weeks. Mice on the high fat diet gained weight faster than their control diet counterparts starting 5-7 weeks, and were glucose intolerant at nine weeks (Figure 1A-D). Nine weeks after diet induction, circulating levels of insulin, c-peptide, and glucose-dependent insulinotropic polypeptide (GIP) were elevated in male mice, but only insulin and c-peptide were elevated in female mice (Figure 1E-J). As shown in clinical models of type 2 diabetes, intraepidermal nerve fiber density was significantly decreased in HFD mice compared to normal chow controls (Supplemental Figure 1). Mechanical paw withdrawal thresholds were measured biweekly to determine the progression of hypersensitivity over time in the HFD mice compared to CON mice, which developed in ten weeks in male mice (Figure 2A) and eight weeks in female mice (Figure 2B). Mechanical thresholds at ten weeks were compared to corresponding fasting blood glucose levels taken within the same week, and no significant correlation between these two factors was found (Figure 2C, D). Paw withdrawal latency to radiant heat or cold were also similar between HFD and CON mice (Supplemental Figure 2A-C).

**Figure 1.**
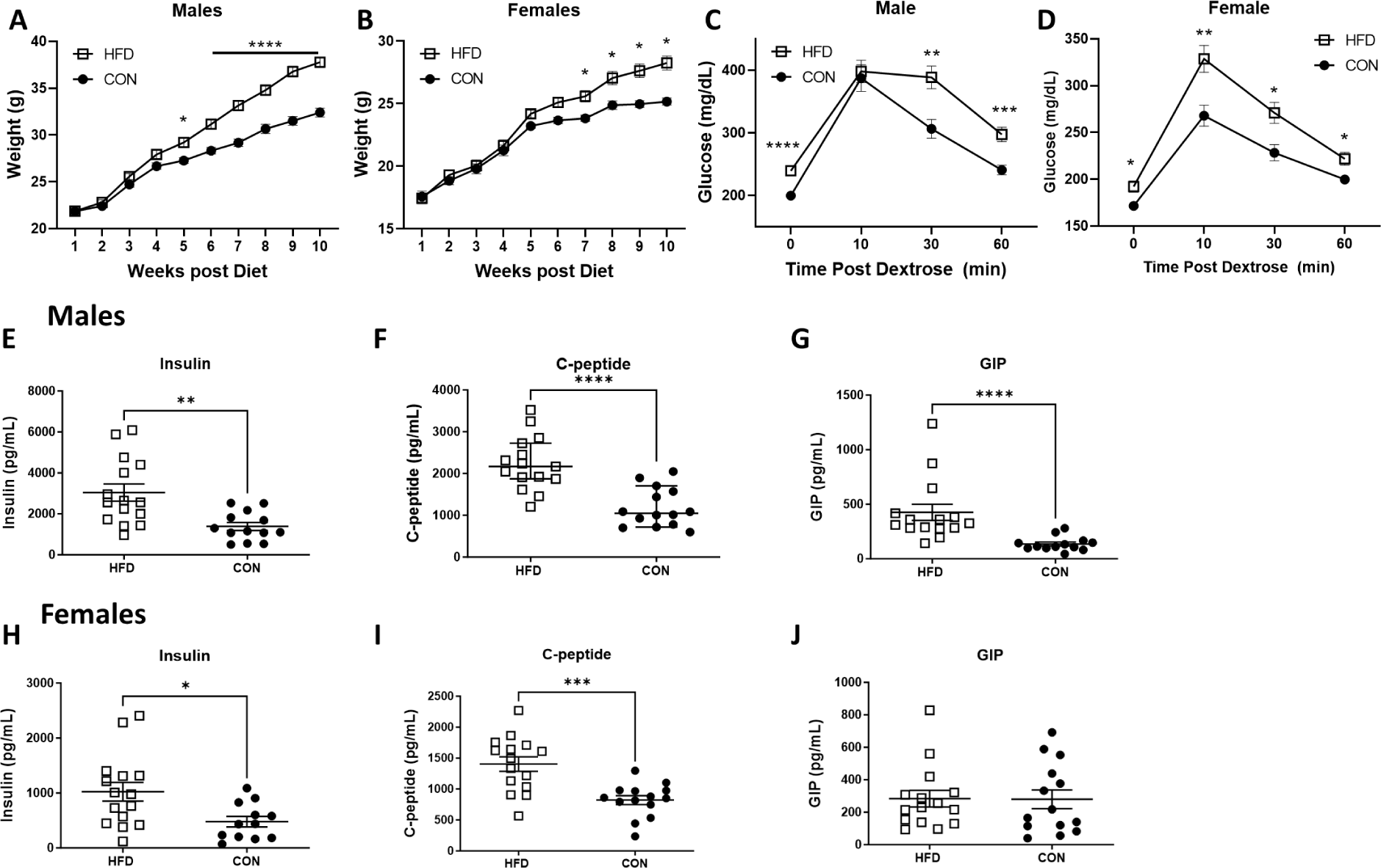
Mice on a high fat diet (HFD) have increased body weight, decreased glucose tolerance, and enhanced levels of serum peptides consistent with type 2 diabetes physiology. Males (A) and females (B) have elevated body weights which is significantly different between HFD and control (CON) diet mice at 5 and 7 weeks after diet induction, respectively (Males: Repeated measures ANOVA, week x diet, F(9,702) = 57.2, p <0.0001; Females: Repeated measures ANOVA, week x diet, F(9,513) = 7.5, p<0.001, Bonferroni post-hoc, n=29-40/group). Glucose tolerance is inhibited in mice fed a HFD at nine weeks post diet induction in male (C) and female (D) mice (Males: Repeated measures ANOVA, week x diet, F(3,234) = 4.3, p=0.006; Females (Repeated measures ANOVA, week x diet, F (3, 174) = 2.836, p = 0.04, Bonferroni post-hoc, n = 30-40/group). Serum levels of insulin (unpaired t-test, p = 0.021), C-peptide (unpaired t-test, p < 0.001) and glucose-dependent insulinotropic polypeptide (GIP, Mann-Whitney U test, p < 0.001) were significantly higher in HFD vs CON diet male mice. In female mice, serum levels of insulin (H) and C-peptide (I) were significantly elevated (unpaired t-test, p = 0.015 and p = 0.0002, respectively), but GIP (J) were similar in HFD compared to CON mice.

**Figure 2.**
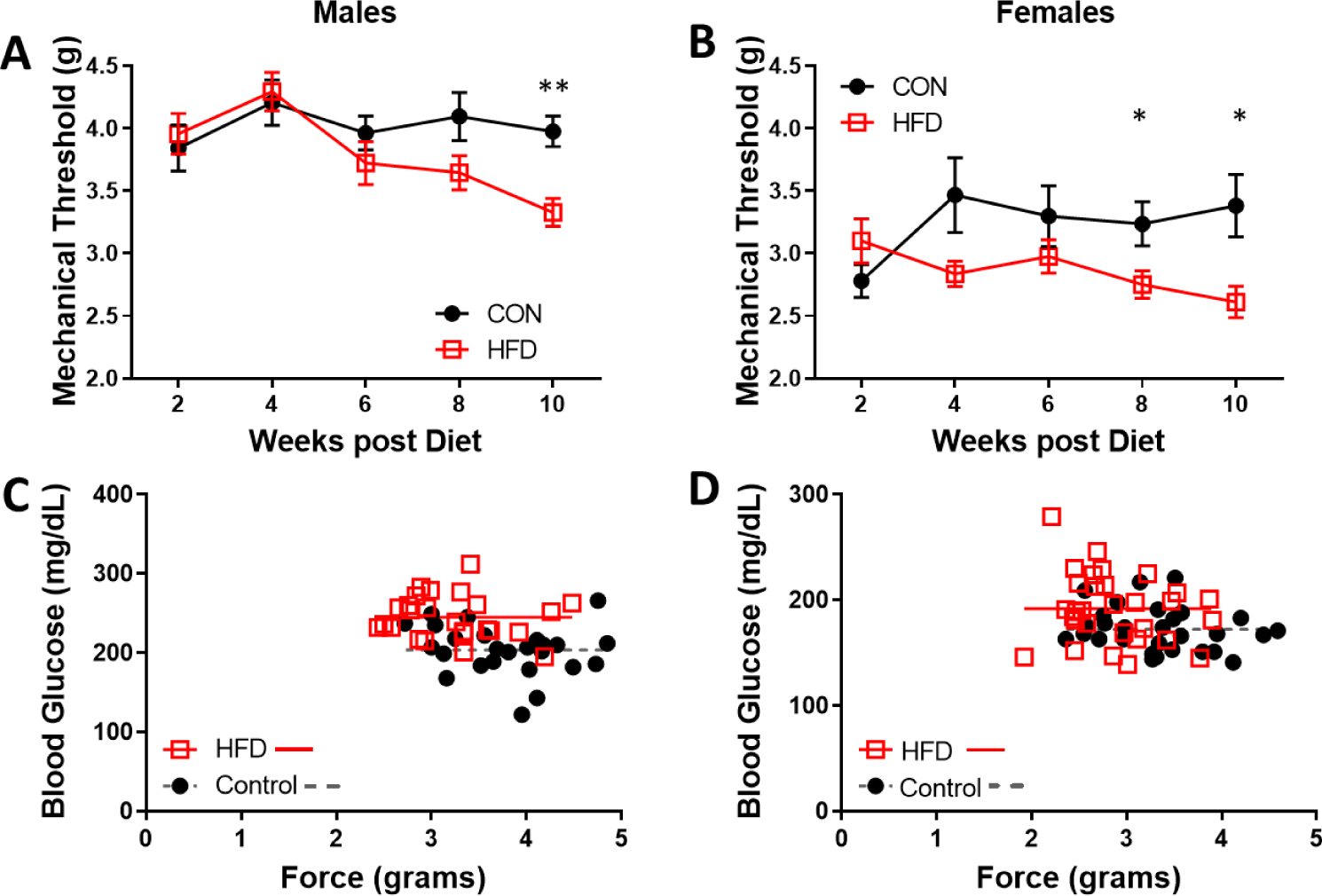
Mechanical paw withdrawal thresholds are inversely correlated with blood glucose levels in mice fed a high fat diet (HFD). (A) Male mice fed a HFD have significantly decreased paw withdrawal thresholds over time compared to mice fed a control diet (CON) (RM ANOVA: F (4, 232) = 2.51, p .043, n=30). (B) Male mice fed a HFD or a CON diet for up to ten weeks do not have a significant correlation between mechanical paw withdrawal thresholds and fasting blood glucose levels (C) Similar decreases in paw withdrawal thresholds were found over time in female mice fed a HFD (RM ANOVA: F (4, 112) = 2.75, p=.032, n =15). (D) There is a lack of correlation between fasting blood glucose and paw withdrawal thresholds ten weeks after either diet introduction in female mice.

### K_ATP_ channel subunit expression in peripheral and central nervous system

The expression of each K_ATP_ channel subunit in the lumbar DRG and spinal cords of HFD and CON mice were measured via qPCR. *Abcc8* was significantly decreased in male HFD mice compared to controls, while Kcnj11 was decreased in both males and females (Figure 3A,B). *Abcc8* and *Kcnj11* were also decreased in the spinal cords of male and female mice (Figure C, D). The expression levels of K_ATP_ channel subunits were correlated with mechanical thresholds in HFD and CON mice. A significant positive correlation was found in the DRG of HFD mice with *Kcnj11* and *Abbc8* expression (Figure 4A, B), but not with *Abcc9* expression (Figure 4C). Expression of Kcnj11 and Kcnj8, were not correlated in the spinal cord with mechanical thresholds, but *Abcc8* and *Abcc9* were positively correlated in HFD mice (Figure 4E-G). Expression levels of KATP channel subunits in CON mice were not significantly correlated in neither DRG nor spinal cord tissues.

**Figure 3.**
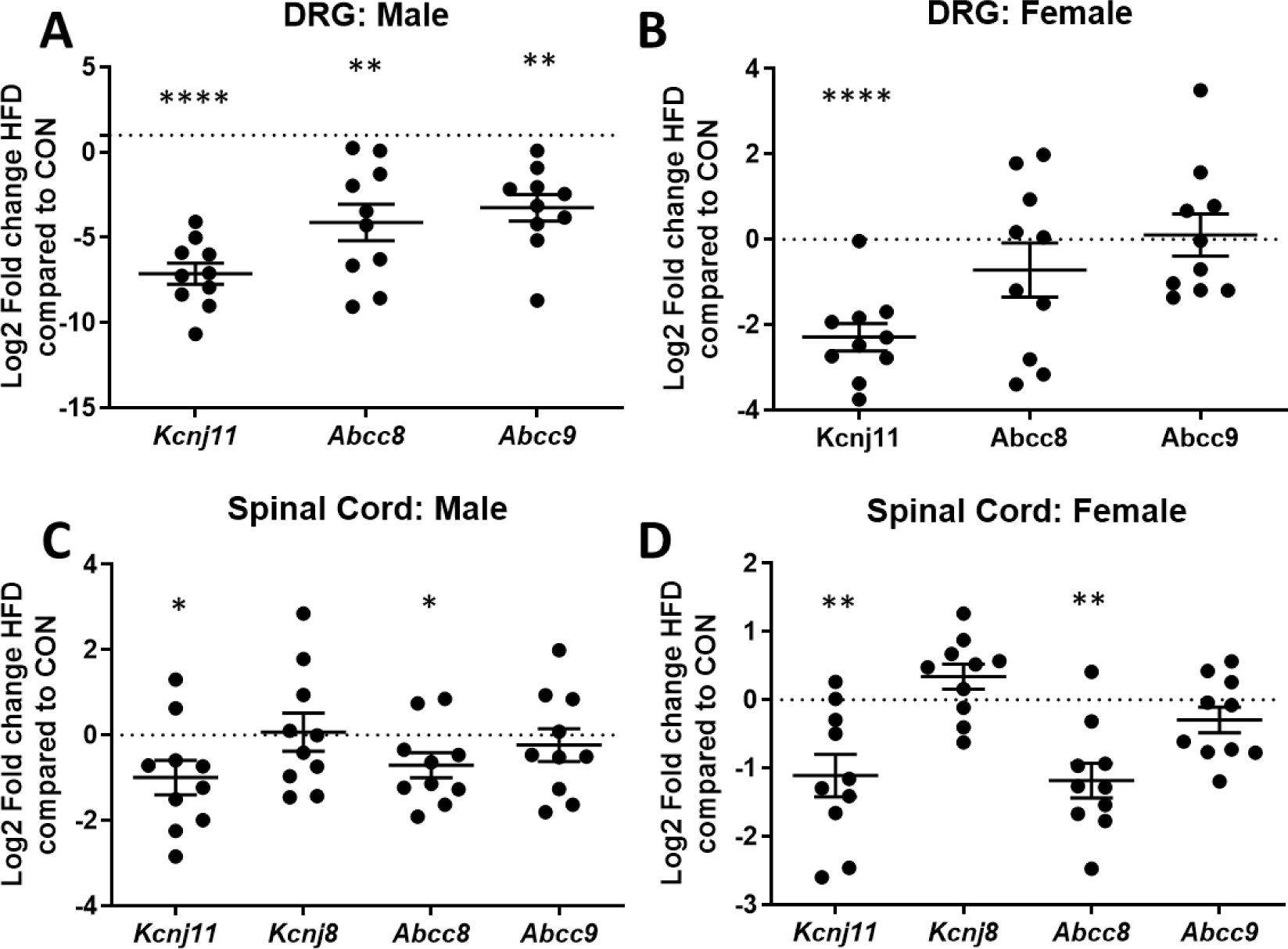
Dorsal root ganglia (DRG) and spinal cord levels of Kir6.2 and SUR1 mRNA are decreased in male and female mice fed a high fat diet (HFD) compared to control diet (CON) fed mice. (A) mRNA levels of the protein products Kir6.2 (Kncj11), SUR1 (Abbc8), SUR2 (Abcc9) are significantly decreased in DRG of male mice on a HFD compared to the average of mice on CON diet (one sample t-test, Kcnj11: t=11.46, p< .0001; Abcc8: t = 3.81, p = .0042, Abcc9: t = 4.17, p = .0024) (B) DRG of female mice on a HFD have significantly decreased mRNA levels of Kcnj11 (one sample t-test, t = 7.043, p < .0001) compared to the average of mice on CON diet. (C) Spinal cord mRNA levels of Kncj11 (one sample t-test, t=2.47, p= .036) and Abbc8 (one sample t-test, t= 2.39, p = .041) are significantly downregulated in male mice and (D) female mice on a HFD compared to the average of mice on CON diet (one sample t-test, Kcnj11: t = 3.57, p = .006; Abcc8: t= 4.65, p=.0012). mRNA levels of Abcc9 or the Kir6.1 gene product (Kcnj8) are not significantly altered for either males nor females in the spinal cord.

**Figure 4.**
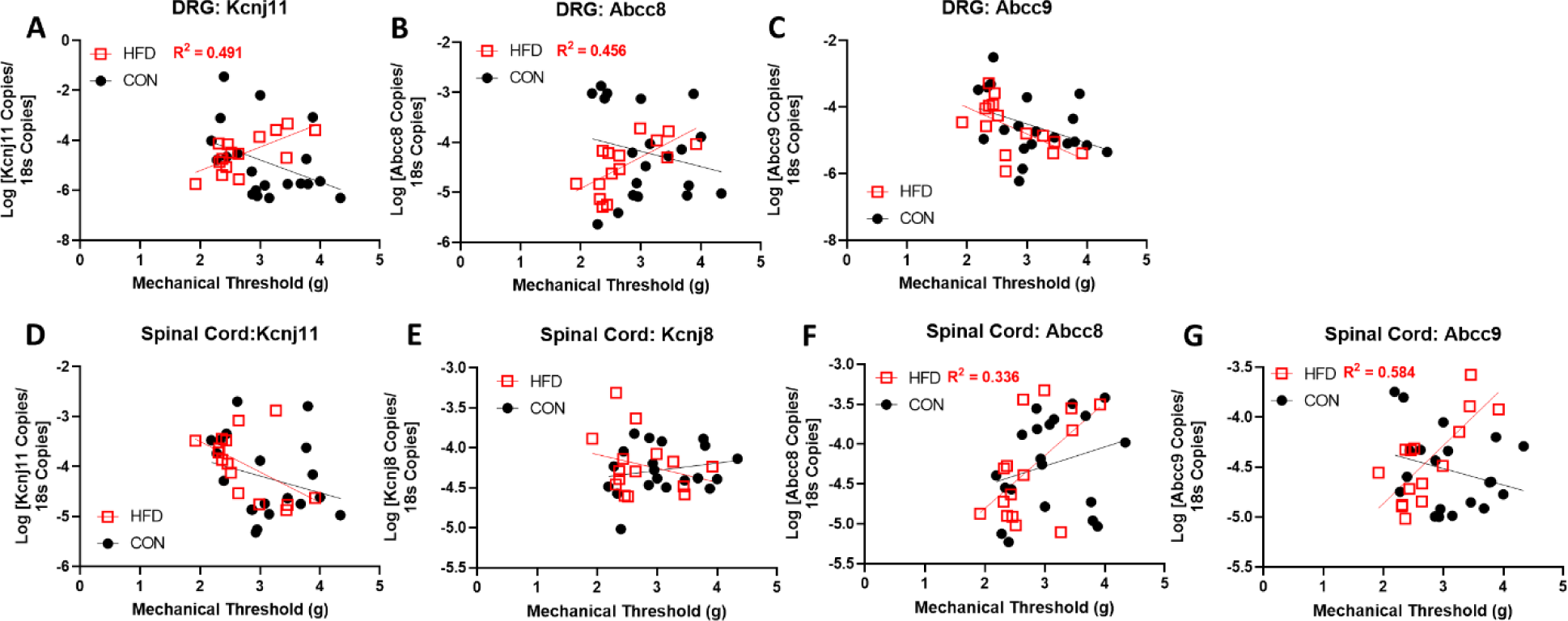
Mechanical paw withdrawal thresholds are positively correlated with Kcnj11 and Abcc8 mRNA expression levels in the DRG (A-C) and spinal cord (D-G) of mice fed a high fat diet (HFD). Mice fed a HFD diet for sixteen weeks have a significant correlation between mechanical paw withdrawal thresholds and number of mRNA copies of (A) Kcnj11 (HFD: Pearson’s correlation coefficient = r(13) = .701, p = .0036) and (B) Abcc8 (HFD: Pearson’s correlation coefficient = r(13) = .675, p = 0.006) in the DRG. (C) There are no significant correlations found between mRNA copies of Abcc9 and mechanical paw withdrawal thresholds in the DRG nor are there significant correlations between (D) Kcnj11 or (E) Kcnj8 mRNA levels in the spinal cord. mRNA levels of (F) Abcc8 (HFD: Pearson’s correlation coefficient: r(13) = .579, p =.024) and (G) Abcc9 (HFD: Pearson’s correlation coefficient = r(13) = .764, p = .0009) are positively correlated with mechanical thresholds in the spinal cord of HFD mice.

### Glyburide/Nateglinide treatment and behavioral testing

Male and female HFD and CON mice were separated into one of three treatment groups 12 weeks after the start of their respective diets. Mechanical thresholds were collected before and after daily administration of glyburide, nateglinide, or vehicle for four weeks. Mechanical thresholds were significantly decreased in HFD diet compared to CON mice treated with vehicle, regardless of systemic treatment (Figure 5A). A cumulative dose response curve indicates that treatment with glyburide or nateglinide, on either CON or HFD significantly decreases the antinociceptive effect of morphine (Figure 5B, C). Several additional tests for general animal wellbeing were also conducted, including nesting and burrowing, which are typically used to assess pain, distress and suffering in rodent models(15). Nesting scores between diet and/or treatment groups were not significantly altered (Figure 6A), but HFD mice had significantly decreased burrowing behavior compared to CON mice (Figure 6B). A significant decrease in motor coordination was also seen in HFD mice compared to CON mice as assessed by time spent on a rotarod apparatus (Figure 6C). These data suggest that HFD mice have higher motor deficits than CON mice, which was corroborated by results of an open field test, where HFD mice traveled less than CON mice (Supplemental Figure 3A). Although significant decrease in mechanical paw withdrawal latencies were seen between diet or diabetes treatments (Figure 5), differences in hyperlocomotor activity were not seen after systemic morphine treatment (Supplemental Figure 3B).

**Figure 5.**
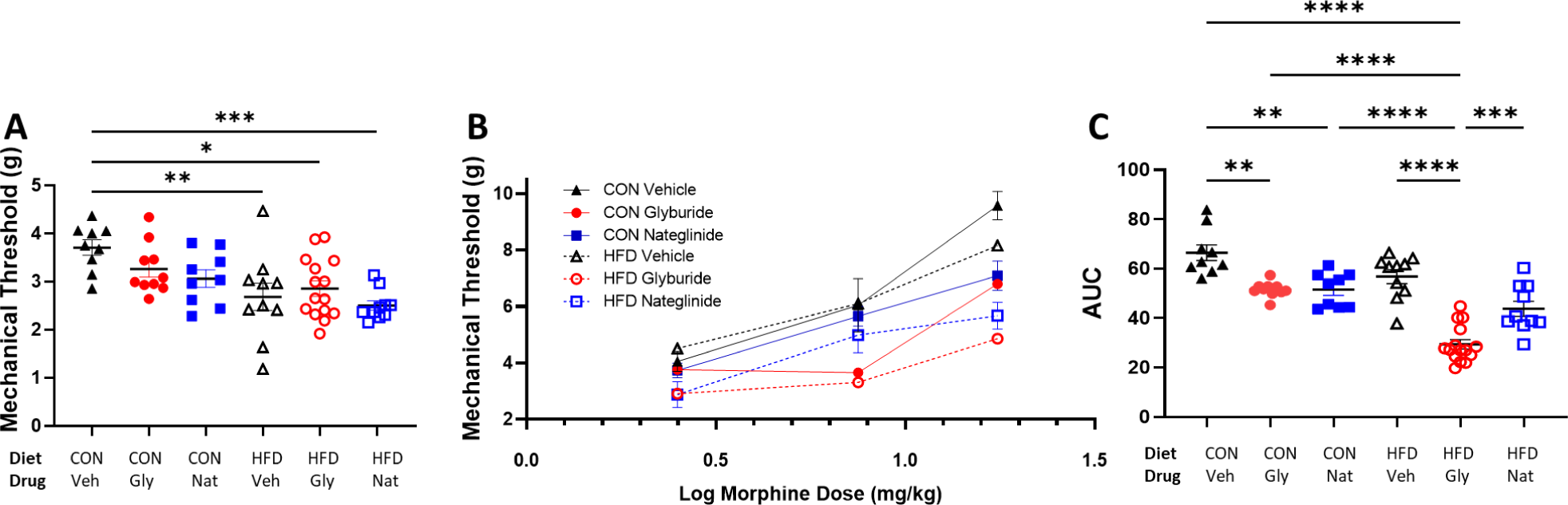
Mechanical paw withdrawal thresholds are significantly decreased in mice fed a high fat diet (HFD). (A) Mice fed a high fat diet for more than three months have decreased mechanical thresholds compared to mice fed a control (CON) diet (F(1, 57) = 18.723, P < 0.001). Dose response curves in control diet and high fat diet mice indicate that glyburide and nateglinide treatment decrease the antinociception of morphine, especially in mice fed a high fat diet. (C) Treatment with glyburide or nateglinide, especially in combination with a high fat diet decreased mechanical thresholds compared to vehicle treatment or control diet. Area under the curve (AUC) of mechanical threshold data obtained in A and B indicate that there is a significant effect of diet (F(1, 57) = 43.5, P < 0.001) and treatment (F(2, 57) = 39.38, P < 0.0001), as well as a significant interaction (diet x treatment, F(2, 57) = 5.53, P = 0.006).

**Figure 6.**
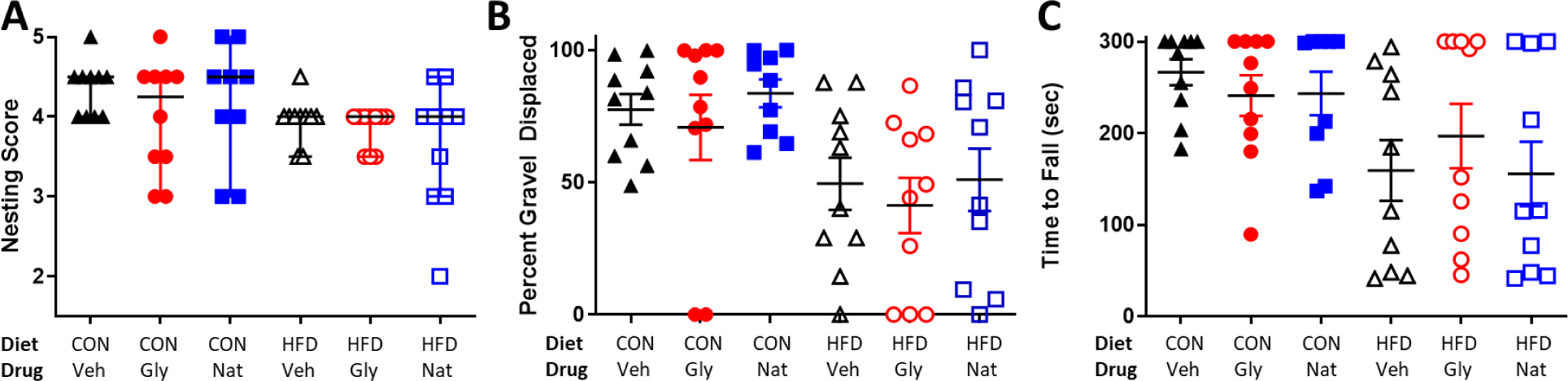
Mice fed a high fat diet (HFD) for over 3 months have decreased burrowing behaviors and latency to falling off a spinning rod. (A) HFD and mice fed a control diet (CON) both exhibited complex nest building behavior and constructed nests regardless of systemic glycemic drug treatment. (B) Mice fed a high fat diet displaced less gravel in a burrowing tube compared to control diet mice (F(1, 53) = 5.088, P = 0.028). (C) The motor coordination, assessed by rotarod test, was significantly decreased in mice fed a high fat diet (F(1, 53) = 7.384, P = 0.009). Daily injections of glyburide (Gly) or nateglinide (Nat) did not significantly impact nesting, burrowing, or motor coordination compared to vehicle (Veh) treated animals. Data in (A) presented as medians with 95% CI. Data in (B) and (C) displayed as means with SEM.

### Upregulation of SUR1 and Kir6.2 and morphine dose response

Since a loss of K_ATP_ channel subunits was correlated with lower mechanical thresholds in HFD mice, a separate cohort of mice were inoculated with an adenovirus in the intrathecal space to upregulate Abcc8 and Kcnj11 in the lumbar spinal cord and DRGs. Abcc8 was significantly elevated in Ad-m-Abcc8 inoculated mice regardless of diet treatment and Kcnj11 was similarly elevated in Ad-m-Kcnj11 inoculated mice (Figure 7A). Mechanical thresholds were higher in CON compared to HFD mice in a cumulative morphine dose response curve (Figure 7B), which was similar to data collected previously (Figure 5B). Expression levels of Abcc8 were significantly correlated with mechanical thresholds, in both HFD and CON mice (Figure 7C), while Kcnj11 mRNA expression was not significantly correlated (Figure 7D). Separate behavioral assays indicate that nesting behaviors were unaltered by either diet nor viral treatment (Figure 8A), but burrowing behaviors were significantly decreased in HFD mice compared to CON mice (Figure 8B). The reduction in burrowing was significantly improved by upregulation of Kcnj11, but not Abcc8 (Figure 8B). Similar to previously collected data, the latency to falling off of a rotating rod was significantly decreased in HFD compared to CON mice, but was not mitigated by viral treatment (Figure 8C).

**Figure 7.**
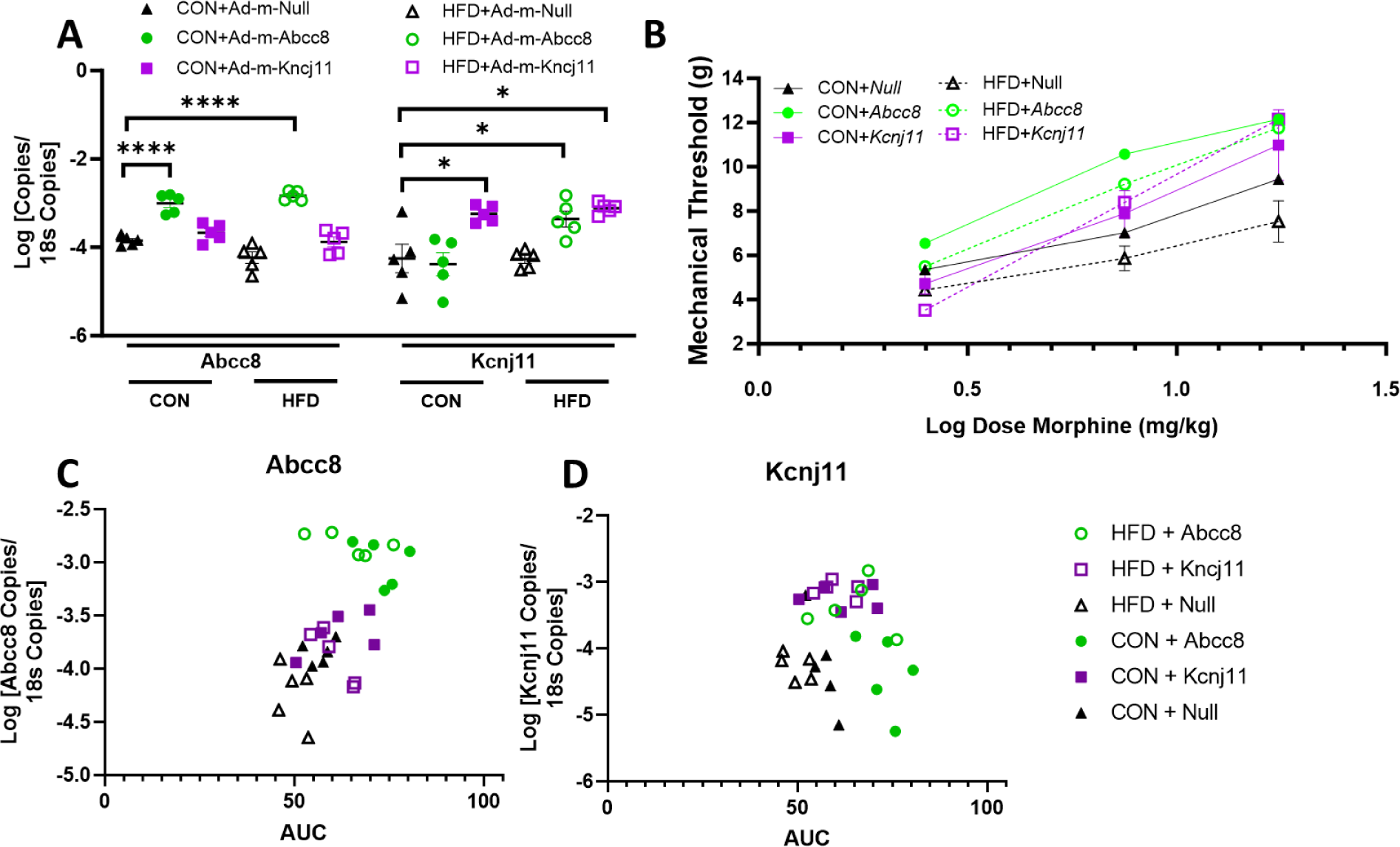
Upregulation of Abcc8 and Kcnj11 increase morphine antinociception in mice fed a high fat diet (HFD). (A) Upregulation of Abcc8 and Kcnj11 mRNA in the spinal cord was detected after intrathecal administration of Ad-m-Abcc8, Ad-m-Kcnj11 compared to a control vector (Ad-m-Null, two-way ANOVA, (F (5, 48) = 17.38, P < 0.0001) (B) Control diet (CON) mice injected with Ad-m-Abcc8 have higher paw withdrawal thresholds to increasing doses to morphine compared to control vector mice (F (2, 12) = 5.861, P = 0.017). (C) Paw withdrawal thresholds in HFD mice are higher in Ad-m-Abcc8 and Ad-m-Kcnj11 mice compared to controls (). (D) Positive correlations exist in HFD and CON mice between the area under the curve (AUC) of paw withdrawals and Abcc8 mRNA expression in the spinal cord (Pearson’s correlation coefficient = r(30) = .589, P = 0.0006)), (E) but not with Kcnj11 mRNA expression (Pearson’s correlation coefficient = r(30) = .00407, P = 0.831).

**Figure 8.**
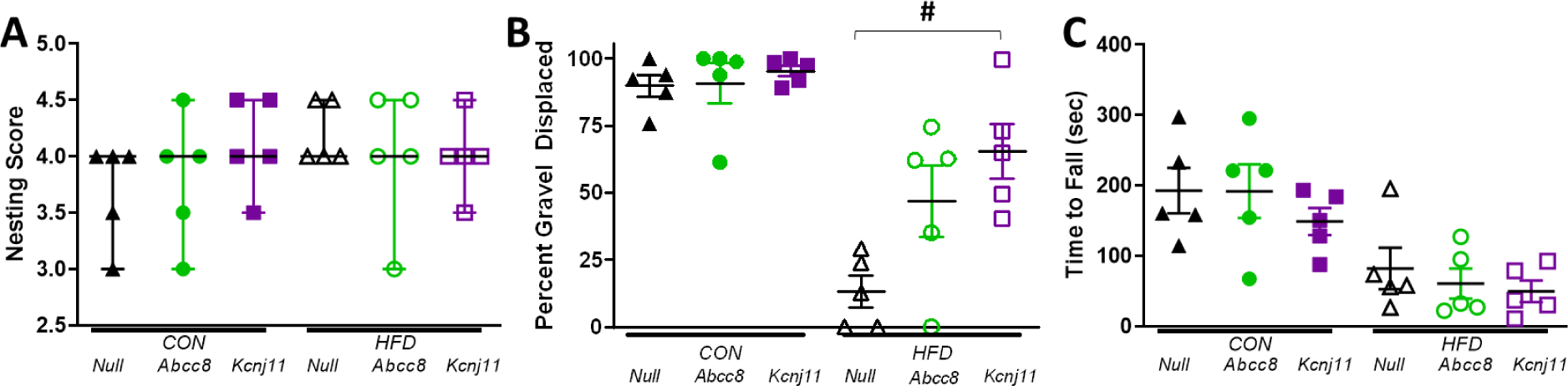
Upregulation of *Abcc8* and *Kcnj11* increases burrowing behaviors in mice fed a high fat diet (HFD) for over 3 months. (A) HFD and mice fed a control diet (CON) both exhibited complex nest building behavior and constructed nests regardless Ad-m-*Abcc8* or Ad-m-*Kcnj11* treatment compared to a control vector (Ad-m-Null). Intrathecal virus injection was performed one week before behavioral testing (10uL, Vector Biolabs) (B) Mice fed a high fat diet displaced less gravel in a burrowing tube compared to control diet mice (F(1, 24) = 57.1, P< 0.001, *). Upregulation of *Kcnj11* also significantly increased burrowing in HFD animals compared to Null controls (F(2, 12) =6.58, P= 0.012, Bonferroni post-hoc, #). (C) The motor coordination, assessed by rotarod test, was significantly decreased in mice fed a high fat diet (F(1, 24) = 26.26, P < 0.001). Data in (A) presented as medians with 95% CI. Data in (B) and displayed as means with SEM.

## Discussion

High concentrations of blood glucose and lipids are considered the primary driver of damage to nerve fibers over time, manifesting in diabetic neuropathy. Diabetic neuropathy, however, is not always associated with epidermal nerve loss(16; 17), nor it is tightly associated with levels of hyperglycemia(18; 19). In this study we did find a significant loss of IENFD in the HFD mice, but we did not find a significant correlation between mechanical paw withdrawal thresholds and fasting blood glucose. Several diabetes models have been used to investigate peripheral neuropathy in rodents, and the diet-induced model containing high fat and carbohydrates was used in our studies in order to most closely emulate physiological and metabolic conditions found in humans. Altered metabolic changes due to hyperglycemia and hyperlipidemia were desired in this study due to the influence of glucose, ATP, and other metabolites on K_ATP_ channel activity and/or expression during diabetes (20–25). Clinical evidence suggests a progressive age-dependent decline in β-cell function during T2DM, including loss of glucose-mediated insulin secretion in the pancreas(26), which is largely dependent on controlled expression and function of K_ATP_ channels(27). mRNA levels for K_ATP_ channel subunits, including *Abcc8* and *Kcnj11* were significantly decreased in both male and female HFD mice in the DRG and spinal cord compared to controls. These data agree with other diabetes studies showing a loss of K_ATP_ channel activity in the central nervous system of rodents (28; 29).

K_ATP_ subunit expression has been found to be altered in several models of chronic pain in rodents and in cellular and rodent *in vivo* diabetes models (30–32), and in human DRG of individuals with painful diabetic neuropathy(31). In vitro, hyperglycemia increases the resting membrane potential of rat DRGs, which is mostly reversed by adding diazoxide, a SUR1-selective agonist(33). Local administration of K_ATP_ channel agonists have been previously shown to reduce hypersensitivity in various rodent models, indicating that activation of K_ATP_ channels in peripheral nerves has either antinociceptive and/or analgesic effects(34–37). Interestingly, the opening probability of K_ATP_ channels typically increases in DRGs after diazoxide application, but this effect is greatly attenuated in rats after spinal nerve injury(38). The loss of K_ATP_ channel function is concurrent with abnormal sensory processing and sensory neuron function, as seen in mice with global loss of Kir6.2 resulting in loss of sensory thresholds and a loss of unmyelinated fibers in the sural nerve(39).

In addition to elevated glucose levels leading to increased intracellular ATP, several metabolic and cellular changes during diabetes could contribute to loss of K_ATP_ channel function and expression. The HFD mice in this study had increased insulin levels, and higher C-peptide and GIP, in addition to lower glucose tolerance compared to CON mice, which are similar to reports from the clinic and previous studies using rodent models (40–43). GIP is one of the key incretin hormones released following ingestion of sugars such as glucose and potentiates glucose-induced insulin secretion from the pancreas in a K_ATP_ channel-dependent manner (44). Incretin-bound receptors such as GIPR increase intracellular cAMP levels through Gs proteins, thereby activating protein kinase A (PKA) and exchange protein activated by cAMP (Epac). Inhibition of adenylyl cyclase or Epac attenuates heat hypersensitivity in db/db mice(45), and increased levels of PKA and/or Epac have been shown to inhibit K_ATP_ channel activity in various cellular models (46; 47). T2DM is also characterized by disproportionate production of intermediary metabolites, including methylglyoxal, that cause cellular damage and increase hypersensitivity and reports of neuropathic pain (48), which has also been shown to inhibit K_ATP_ channel expression over time (49).

Sulfonylureas, such as glyburide, haven been shown to inhibit antinociception in several animal models, particularly after opioid administration (36; 50; 51). Since K_ATP_ channels are downstream of opioid receptor signaling, it would make sense that inhibition of potassium currents would decrease the hyperpolarizing effects of opioids. The antinociceptive effect of morphine was significantly decreased in HFD mice compared to CON mice, and this was further exacerbated in mice treated with glyburide or nateglinide. Nerve conduction measures are typically used to determine neuropathy disease severity in the clinic(52), and we did find small changes in conduction velocity or amplitude in HFD compared to CON mice. Treatment with either diabetes medication did not impact nerve conduction nor other behavioral measures (e.g. nesting, burrowing, open field mobility). This indicates that the HFD mouse model potentially affected small and large nerve fibers involved in sensation and locomotion, which is typical for most diet-induced T2DM models in rodents(53). Diet and drug treatment did impact morphine-induced antinociception, but did not have an effect on morphine-induced hyperlocomotion.

Possible explanations include the low penetration of sulfonylureas or meglitinides across the neurovascular unit, and that hyperglycemia and hyperlipidemia do not impact all neurons in the brain equally. In addition to morphine, we also demonstrated an antinociceptive effect of pregabalin, which was similar across CON and HFD mice. Interestingly, the anticonvulsant gabapentin did not have a significant effect on mechanical paw withdrawal thresholds across a wide range of doses tested, indicating that gabapentin may have limited analgesic efficacy, similar to reports from the clinic(54).

Significant positive correlations between mechanical paw withdrawal thresholds and K_ATP_ channel subunit expression were found in DRG and spinal cord of diabetic animals. In the peripheral nervous system, we found correlation between Kcnj11 and Abcc8, and a correlation between *Abbc8* and *Abcc9* in the spinal cord. We then employed a viral strategy to upregulate *Abcc8* and *Kcnj11* in the spinal cord of CON and HFD mice to determine if restoration of K_ATP_ channel expression could improve morphine antinociception. Interestingly, *Abbc8* upregulation did improve the mechanical paw withdrawal thresholds, particularly in HFD mice whereas *Kcnj11* had a much lesser effect. In vitro studies indicate that SUR1 is more sensitive to changes in ADP/ATP levels than SUR2 when coupled to Kir6.2 (55) indicating that the sulfonylurea units are key players in metabolic coupling to resting membrane potentials. SUR1 and SUR2 are the molecular targets for several diabetes medications (e.g. glyburide, gliplizide, repanglinide), rather than the Kir subunits (56) and also the target for ATP/ADP which is a key parameter to regulating cellular metabolism. Upregulation of SUR1 enhanced morphine antinociception especially in HFD mice, possibly due to changes in metabolites, energy expenditure, and mitochondrial function during diabetes that may impact sulfonylurea subunits more greatly than Kir6.2 functioning.

K_ATP_ channels are expressed in several organs/tissues including blood vessels (Kir6.1/SUR2) and ventricular myocytes (Kir6.2/SUR2). Although pancreatic and neuronal KATP channel subunits are typically Kir6.2/SUR1, several secretagogues targeting K_ATP_ channels do not selectively target one subunit over another. Several early studies have suggested an associated risk between sulfonylurea use and adverse cardiovascular events, particularly infarction and stroke, but recent studies have had mixed results(57–61). Several possibilities exist for the inconsistencies in determining trends, including duration and severity of diabetes, choice of reference group, previous history of cardiovascular events, and length of follow-up. One important consideration is the type of sulfonylurea used by the patient and the differences in formulation (e.g. extended release) and differences in pharmacological properties among these drugs, could result in different levels of cardiovascular risk because of differences of direct action on vessels/cardiomyocytes or glycemic levels indirectly affecting heart function(62). The results of these studies suggest that diabetics using drugs targeting K_ATP_ channels could have altered sensory perception and lower efficacy of opioid medications to reduce pain. To date, we are unaware of any large-scale cross-sectional analyses of data to characterize trends in the use of sulfonylurea/metaglinide drug types among patients taking opioids versus non-opioid medication for chronic pain. Given the issues previously raised with studies investigating cardiovascular adverse events and sulfonylureas, it would be useful to continue these efforts with neuropathy and opioid use in diabetics, with careful controls for any potential confounding of data.

Chronic administration of morphine in mice decreases the affinity of K_ATP_ channels for glyburide in the spinal cord, suggesting that compensatory changes in neurons leads to a decrease in potassium efflux and less hyperpolarization, resulting in less antinociception or analgesia(63). Coupled with data suggesting that K^+^_ATP_ channels are involved in DRG hypersensitivity during hyperglycemia, this would suggest that treatment of diabetes with sulfonylurea/metaglinide drug types coupled with chronic opioid use for would be problematic for patients with diabetic neuropathy(64). Of course, alternative diabetic medications exist beyond sulfonylureas. Currently, metformin is the most prescribed antidiabetic drug (∼50% of type 2 diabetes patients) which has remained consistent over the past ten years(65). The mechanism of action of metformin is probably multifactorial, as it is reported to increase the effects of insulin and also suppress endogenous glucose production from the liver by stimulating the activation of phosphodiesterase (PDE) and downregulation of cAMP signaling. Metformin also increases cellular AMP to inhibit adenylyl cyclase (AC) and thus suppresses cAMP synthesis. Glucagon-like peptide 1 (GLP-1) agonists are a class of drugs with ever increasing popularity for their ability to stimulate insulin secretion. GLP-1 also enhances cAMP production to facilitate a PKA-induced closure of K_ATP_ channels(47). Whether or not metformin or GLP-1 agonists directly or indirectly influences ion channel excitability, and therefore influences incidence of chronic pain, particularly from diabetes is up for debate(66–68). According to a recent cohort study, patients with diabetes who taking opioids were at increased risk of early mortality compared to diabetic and non-diabetic participants who did not use opiates and non-diabetics using opiates(69). Further investigation on opioid and anti-diabetic drug combinations a relevant future area of study. Further studies are also needed to verify whether the long-term use of the antidiabetic drugs increase the susceptibility of patients to neuropathy and/or loss of analgesic efficacy due to downstream targeting of ion channels.

## Acknowledgements

This work was supported through a University of Minnesota Academic Health Center Faculty Development Grant to AHK and MLG and R01DA051876 awarded to AHK. The authors do not report any conflicts of interest.

---

**Supplemental Figure 1.**
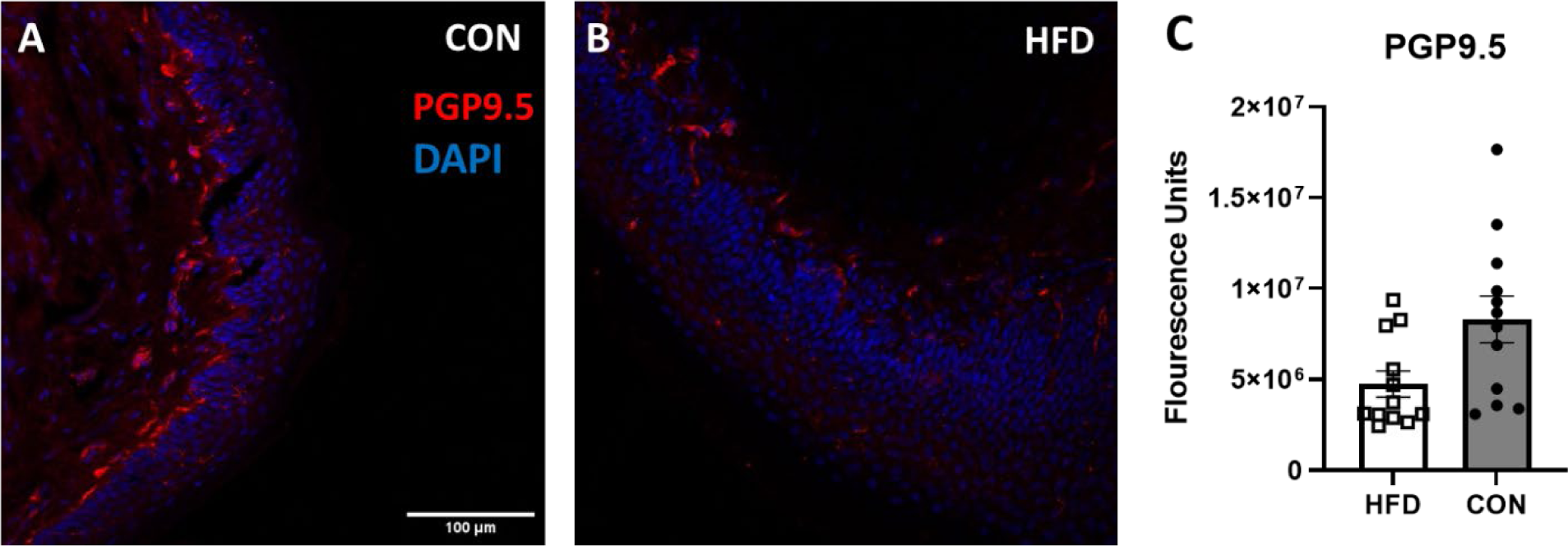
Reduction in intraepidermal nerve fiber density (IENFD) in mice fed a high fat diet (HFD) for 4 months. (A) Photomicrographs of intraepidermal nerve fiber fibers in control diet (CON) or (B) high fat diet (HFD) for sixteen weeks. Hindpaw skin was sectioned and stained with PGP9.5, DAPI staining indicates epidermal layer of skin. Scalebar = 100 uM. (C) Flourescence intensity of PGP9.5 was significantly reduced in HFD compared to CON mice (unpaired t-test, P = 0.0236).

**Supplemental Figure 2.**
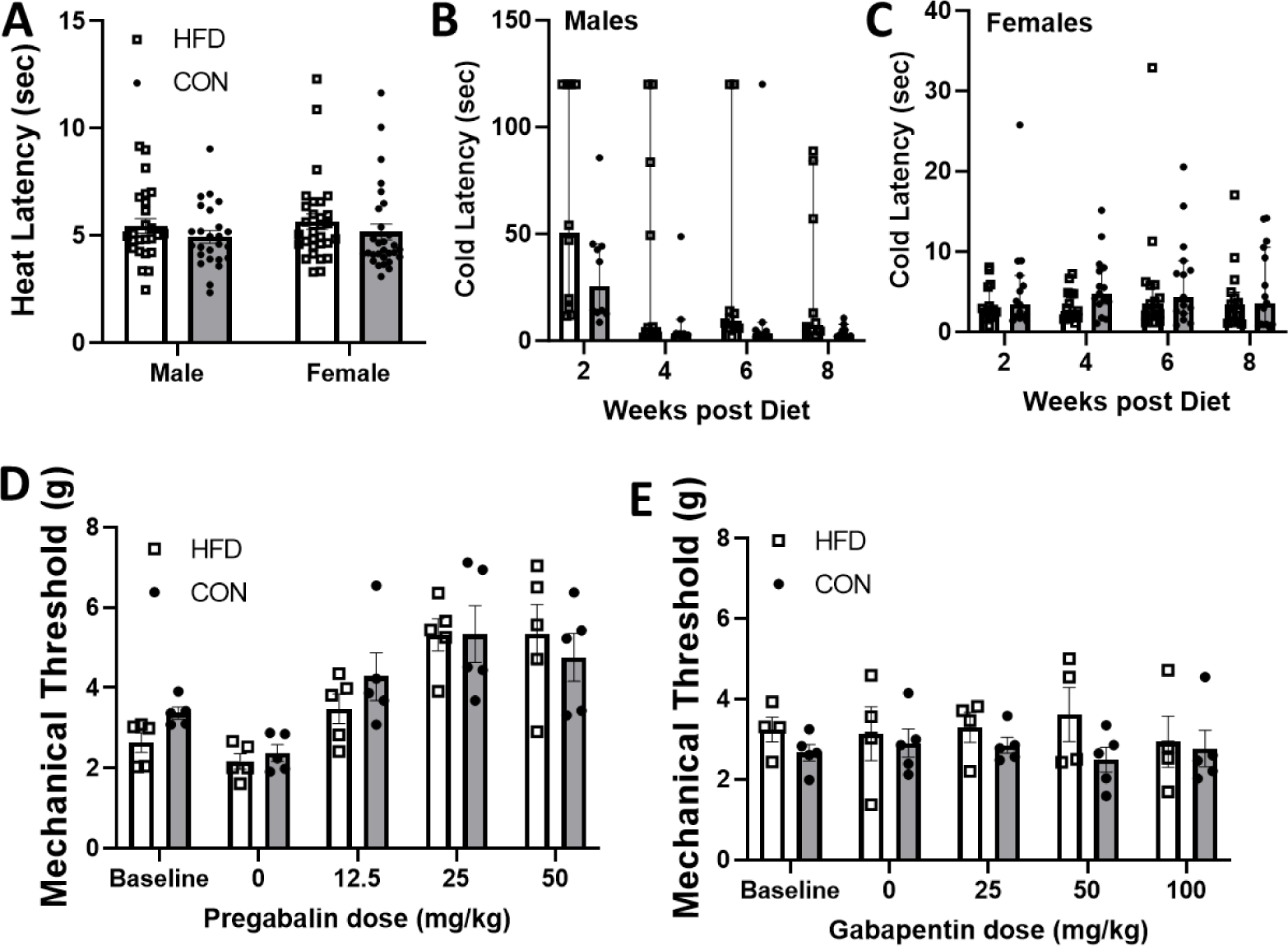
Lack of changes to heat and cold thermal thresholds and pregabalin and gabapentin antinociception in mice fed a high fat diet (HFD). (A) Hindpaw thermal thresholds to radiant heat after 10 weeks and cold latency to an aluminum block kept on ice are not significantly altered in (B) male or (C) female HFD mice compared to mice fed a control diet (CON). (D) Mechanical withdrawal latencies are not significantly impacted by diet status after systemic administration of pregabalin (0-50 mg/kg, sc) or (E) gabapentin (0-100 mg/kg, sc). Data in A, D, and E presented as mean ± SEM, data in B and C presented as median ± 95% CI.

**Supplemental Figure 3.**
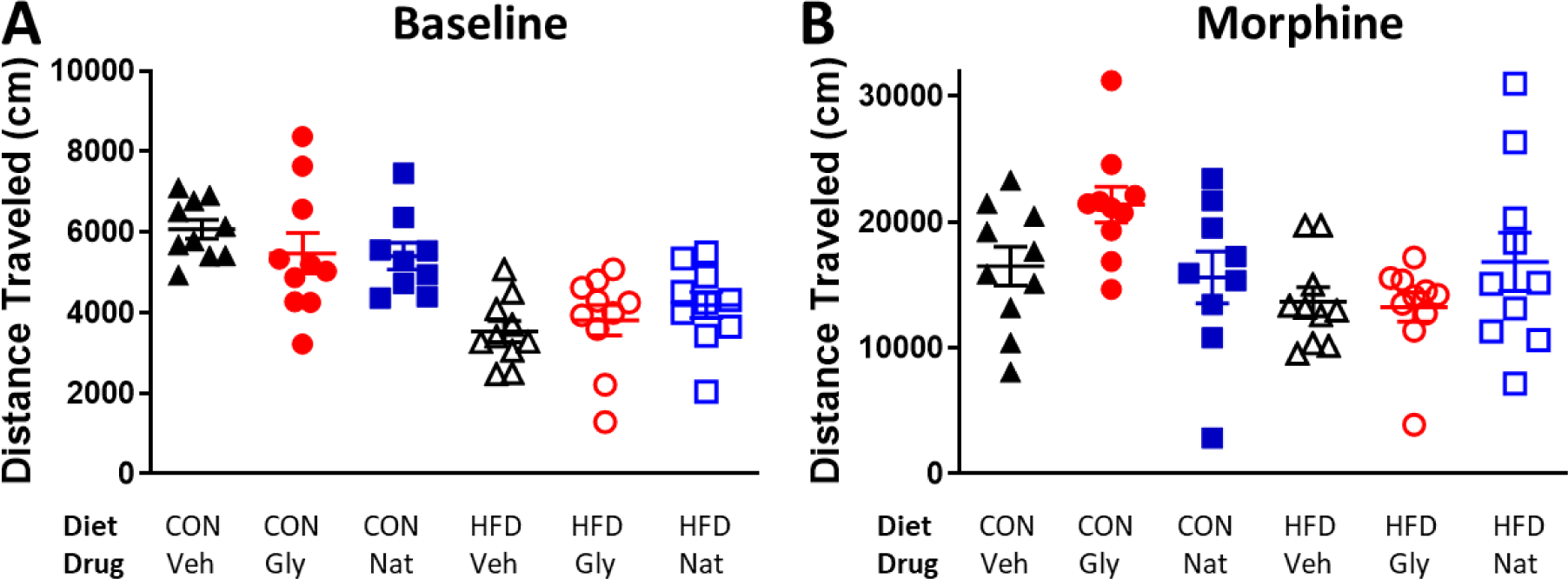
Decreased baseline locomotor activity in an open field behavior test in mice fed a high fat diet (HFD) for over 3 months. (A) Mice fed a control (CON) diet traveled further during a 15-minute baseline period regardless of systemic glycemic drug treatment compared to HFD mice (ANOVA: F(1,48) = 33.29, P< 0.001). (B) Hyperlocomotion induced by morphine (15 mg/kg, sc, 30 minutes) was not significantly affected by diet or glycemic drug treatment.

**Supplemental Figure 4.**
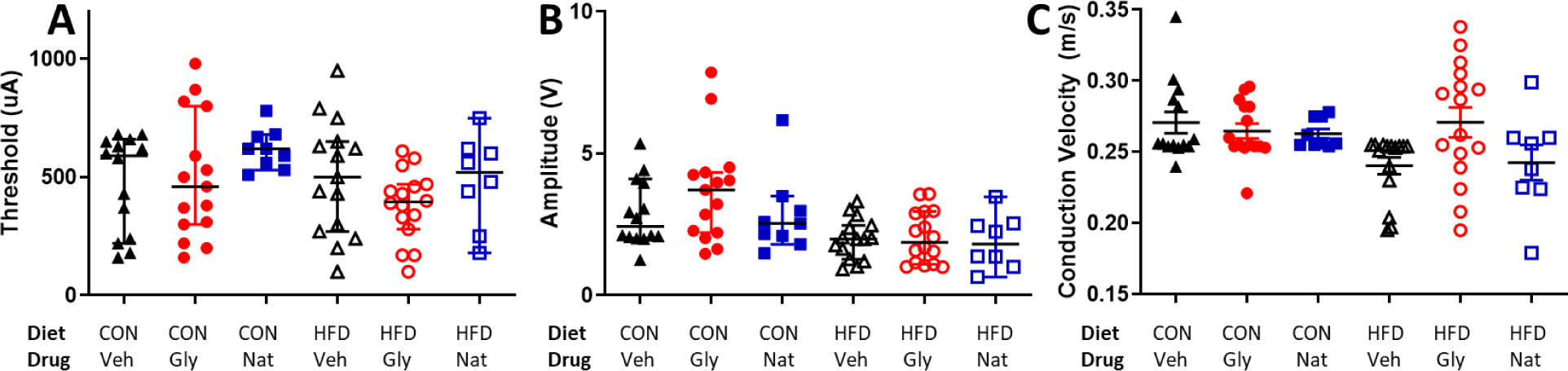
Mice fed a high fat diet (HFD) for over 3 months have altered C-fiber compound action potential amplitudes and conduction velocity. (A) HFD and mice fed a control diet (CON) have similar stimulation thresholds regardless of systemic glycemic drug treatment of either glyburide (Gly) or nateglinide (Nat) compared to vehicle treatment (Veh). (B) Mice fed a high fat diet have lower maximum amplitudes (F(1, 71) = 15.7, P < 0.001). (C) Conduction velocity of C-CAPs were significantly decreased in mice fed a high fat diet (F(1, 71) = 4.20, P = 0.044). Data in (A) presented as medians with 95% CI. Data in (B) and (C) displayed as means with SEM.

## Notes

### Competing Interest Statement

The authors have declared no competing interest.

